# Structural Modeling of Peptide Toxin - Ion Channel Interactions using RosettaDock

**DOI:** 10.1101/2022.06.29.498146

**Authors:** Diego Lopez Mateos, Vladimir Yarov-Yarovoy

## Abstract

**SUMMARY:** Voltage-gated ion channels play essential physiological roles in action potential generation and propagation. Peptidic toxins from animal venoms target ion channels and provide useful scaffolds for the rational design of novel channel modulators with enhanced potency and subtype selectivity. Despite recent progress in obtaining experimental structures of peptide toxin – ion channel complexes, structural determination of peptide toxins bound to ion channels in physiologically important states remains challenging. Here we describe an application of RosettaDock approach to structural modeling of peptide toxins interactions with ion channels. We tested this approach on 10 structures of peptide toxin-ion channel complexes and demonstrated that it can sample near-native structures in all tested cases. Our approach will be useful for improving understanding of the molecular mechanism of natural peptide toxin modulation of ion channel gating and for the structural modeling of novel peptide-based ion channel modulators.

## INTRODUCTION

Voltage-gated ion channels (VGICs) are transmembrane proteins that enable ions to permeate through the cell membrane in response to membrane depolarization. VGICs are involved in diverse and crucial physiological processes such as transmitting the nerve impulse or muscle contraction (Catterall, 1986, 2010). Mammalian voltage-gated sodium (Na_V_) and voltage-gated calcium (Ca_V_) channels share a common architecture with a core α-subunit and a set of auxiliary subunits (Catterall, 2000; Pan et al., 2018). The α subunit folds into four homologous tandem domains, each comprising six transmembrane segments (S1-S6). The first four transmembrane segments (S1 to S4) form the voltage-sensing domain (VSD), while the last two transmembrane segments (S5 and S6) form the pore domain (PD). The four VSDs are located in the periphery of the channel, while the PDs form the central ion permeation pathway. S4 segments in each VSD contain the gating charges and move up (activated state) or down (deactivated or resting state) in response to changes in membrane potential, which provides the mechanical work required to open or close the channel (Catterall, 2010). Voltage-gated potassium (K_V_) channels present a similar architecture, but each domain comprises an independent polypeptide chain resulting in homo or heterotetrameric structures (Attali et al., 2021; Yellen, 2002).

Due to their physiological relevance, VGICs are important molecular targets for treating a large diversity of diseases (Garcia and Kaczorowski, 2021; Wulff et al., 2019). Achieving high subtype selectivity for molecules targeting specific VGICs is of primary importance to not impair crucial physiological functions mediated by off-target channels. Pore blockers such as tetrodotoxin (TTX) and 4-aminopyridine (4-AP) can respectively inhibit currents mediated by many Na_V_ and K_V_ channels (Catterall, 1986; Rodríguez-Rangel et al., 2020). However, the high degree of sequence conservation of the pore domain across different ion channels hinders accomplishing subtype selectivity by targeting this region. Peptidic gating modifier toxins (GMTs) from animal venoms target ion channel VSDs and are promising scaffolds for novel peptide design with a large subtype selectivity (Herzig et al., 2020; Wulff et al., 2019). By binding to the VSDs, these venom peptides can trap the channel in a functional state and alter its gating and dynamic properties (Xiao et al., 2010).

Obtaining experimental structures of peptide toxin – ion channel complexes is crucial to understanding the peptide toxin binding mechanisms and the molecular basis of their channel modulation and, ultimately, to design new peptides with enhanced potency, specificity, and stability to be useful as molecular tools to study ion channel activity and as novel therapeutics to treat ion channel-dependent diseases. However, several technical difficulties have hampered the structural determination of peptide toxins bound to human VGICs in different states. Firstly, purification and crystallization of human Na_V_ and Ca_V_ channels present an important experimental challenge due to the large size of these proteins (∼250 kDa) and the importance of keeping the channels in a membrane-like environment. Secondly, VGICs adopt multiple states and associated conformations induced by changes in the membrane potential *in vivo*, which is difficult to replicate during the purification and structural determination process.

In recent years, impressive progress has been made in resolving experimental structures of venom peptides bound to ion channels in multiple states (Banerjee et al., 2013; Clairfeuille et al., 2019; Gao et al., 2021; Jiang et al., 2021a; Pan et al., 2019; Shen et al., 2018, 2019; Wisedchaisri et al., 2020; Xu et al., 2019; Yu et al., 2005) (Figure 1).

**Figure 1.**
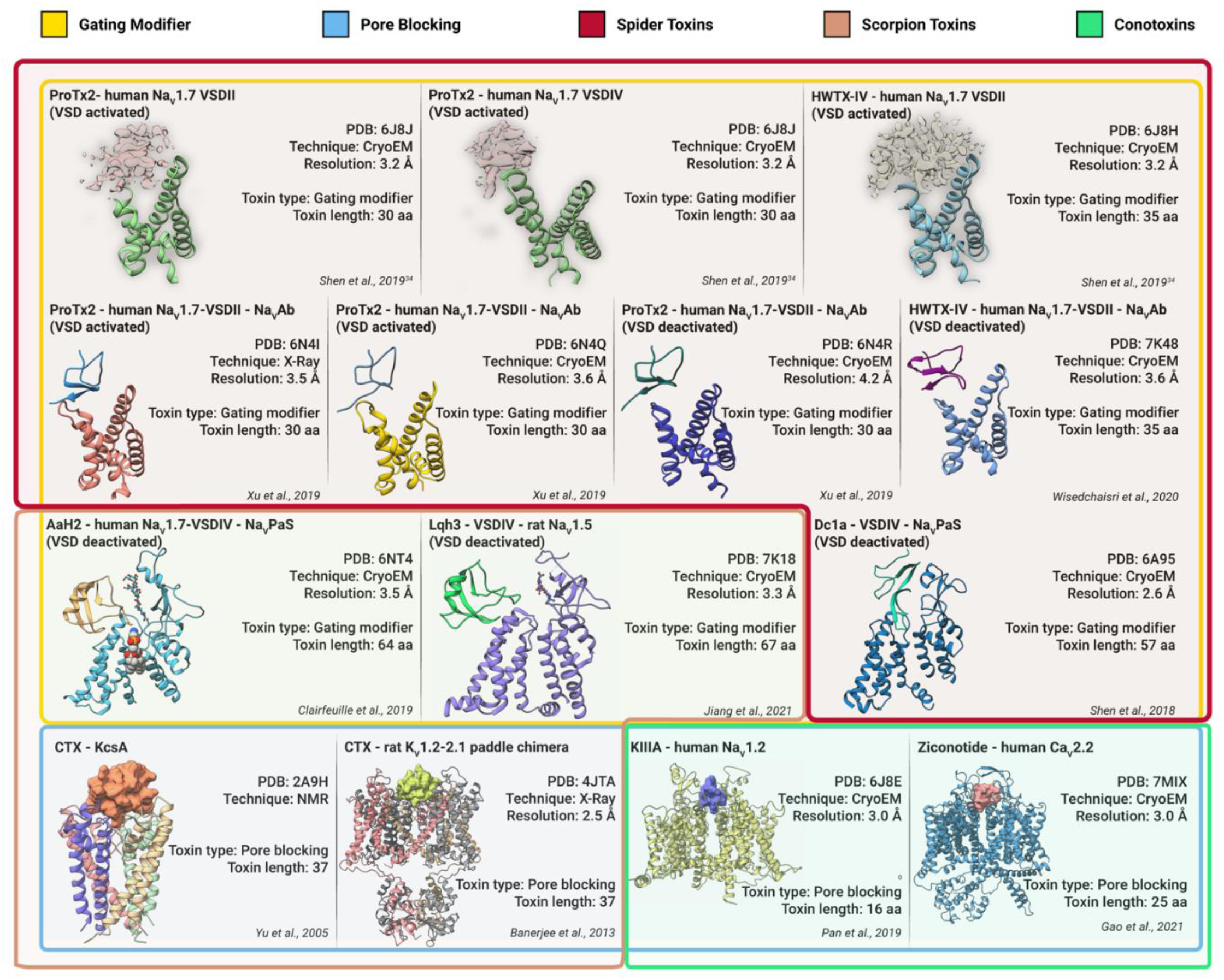
Native structures of peptide toxin – ion channel complexes. Available native structures of peptide toxin – ion channel complexes at the Protein Data Bank (PDB) are shown. Colored lines around the structures group them according to two criteria: type of toxin (gating modifier or pore blocking) and animal source (spider, scorpion, or cone snail). For structures only involving interactions at the VSD, only this domain is shown (6J8J, 6J8H, 6N4I, 6N4Q, 6N4R & 7K48). 6J8J and 6J8H cryoEM structures did not have enough resolution to build atomic models of the toxin, therefore, only the toxin EM densities are shown. The PD is shown for structures involving GMTs that interact with this region (6NT4, 7K18 & 6A95); glycans and lipids are highlighted using sticks and sphere representations respectively. For pore blocking toxins, the whole corresponding channel is shown, and toxins are highlighted in surface representation.

The therapeutic potential of animal toxins has already been harnessed (Bordon et al., 2020; Muttenthaler et al., 2021; Pennington et al., 2018), resulting in unique treatments like ziconotide for the treatment of severe chronic pain (McGivern, 2007) or the lizard-derived toxin exenatide for the treatment of treat type 2 diabetes (Holz and Chepurny, 2003). However, enhancement of the native peptide toxins to better fulfill their therapeutic effects has proven difficult and expensive (Neff and Wickenden, 2021). When atomistic peptide – ion channel structures are available, structure-guided computational design can be used to improve the binding features of the peptide, boosting the drug development process (Nguyen and Yarov-Yarovoy, 2022). There is an immense diversity of venom peptides targeting ion channels (Herzig et al., 2020; Hung et al., 2017), and depending on resolving experimental structures of relevant toxins bound to their targets poses a bottleneck for our opportunities to exploit their potential for pre-clinical optimization and future clinical applications. We propose that computational tools can be used to predict and model the interactions between peptide toxins and ion channels with high accuracy, offering an alternative approach to acquire the structural data required to rationally design novel ion channel modulators and understand the molecular mechanism of peptide toxin action.

Using the Rosetta software modeling suite (Bender et al., 2016; Leman et al., 2020; Rohl et al., 2004), we developed a robust protocol to model peptide – ion channel interactions and tested its performance on ten available experimental structures of venom peptide – ion channel complexes. Rosetta sampled near-native conformations in all tested cases. Incorporating Rosetta membrane energy function in cases with membrane-embedded peptide toxins resulted in enrichment of near-native models and improved scoring compared to the standard Rosetta energy function. Pore-blocking conotoxins required biased sampling to generate near-native models within a narrow and constrained binding site. Identification of near-native models was accomplishable using selection criteria based on the model’s total and interface scores. Importantly, we observed that the explicit presence of N-glycosylation of the channel, when located at the interface with the toxins, might be necessary for scoring in cases involving α-scorpion toxins. Our results provide a novel approach and point of reference to model and explore peptide toxin – ion channel interactions. We anticipate that computational modeling and design of peptide – ion channel interactions will play a vital role in the drug development process targeting ion channels as computational structural biology approaches become increasingly accurate.

## RESULTS

### Modeling protocol and test cases

We applied the RosettaDock protein-protein docking protocol (Chaudhury et al., 2011) to model the available structures of peptide toxin – ion channel complexes (Figure 1). The RosettaDock protocol is a multi-scale Monte-Carlo based algorithm in which the peptide is randomly perturbed starting from initial positions near the channel in terms of translation, rotation, backbone torsional angles, and side-chain rotamers (Chaudhury et al., 2011). We performed an exhaustive sampling of peptide toxin – ion channel complexes to identify near-native models using Rosetta scoring.

Our modeling protocol workflow comprised of the following steps (see Figure 2 and Methods): 1) optimization of the channel and peptide structures within the Rosetta framework (fast relax and EM density refinement) (Conway et al., 2014; Wang et al., 2016); 2) transformation into membrane coordinates to use a membrane energy function (franklin2019) (Alford et al., 2020) and determination of membrane geometry parameters and transmembrane spans; 3) initial manual placement of the peptide in different positions around the putative binding site with each location providing a different input for the subsequent steps; 4) prepacking to optimize side-chains outside the docking interface (Chaudhury et al., 2011); 5) RosettaDock application (Chaudhury et al., 2011); 6) Analysis of the generated models. The information contained in the scoring of the models is used to identify near-native models.

**Figure 2.**
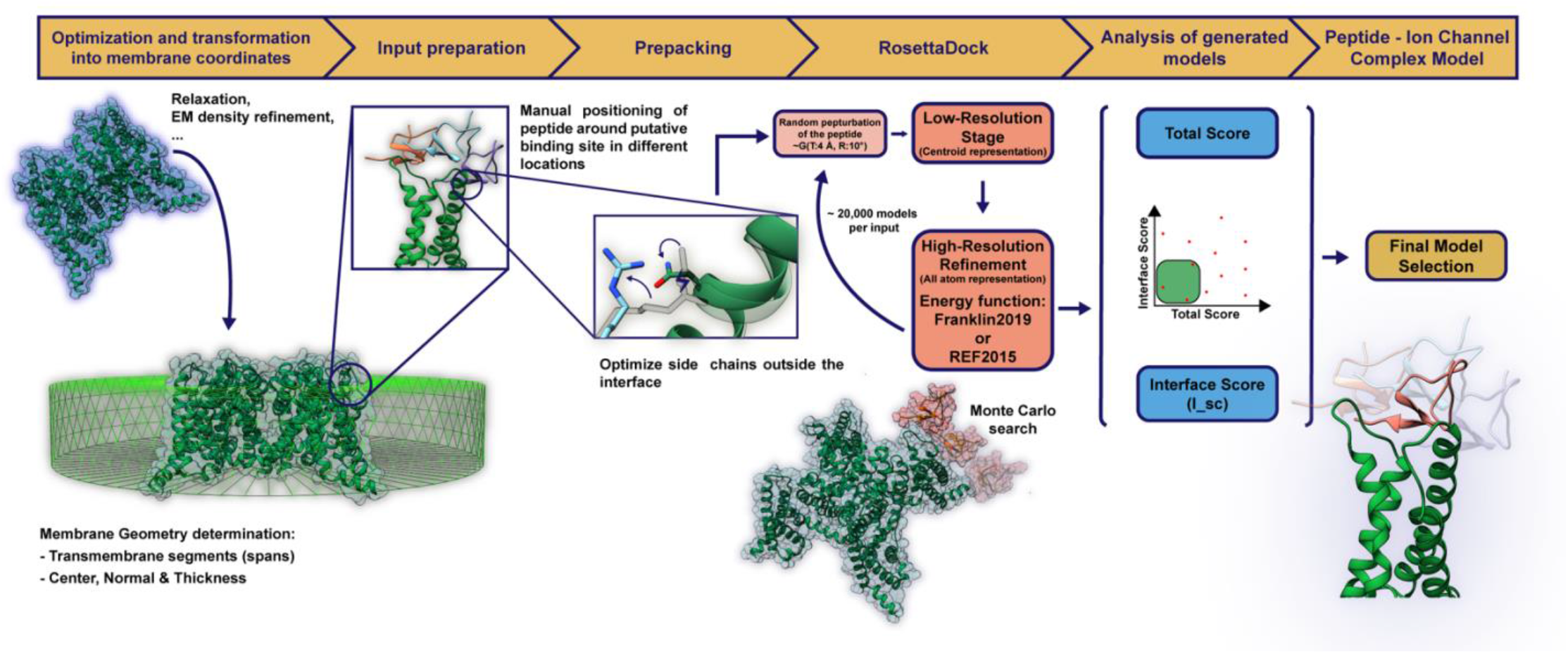
Workflow of protein-protein docking protocol to model the interactions between peptide toxins and ion channels. Sequential steps in the modeling protocol are indicated in the yellow bar on the top. Below, visual illustration and details of each step are shown. Further details, code and execution options have been provided in Methods.

We tested the latest Rosetta membrane energy function (franklin2019) (Alford et al., 2020) and compared its performance with the standard Rosetta energy function (REF2015) (Alford et al., 2017) for peptide toxin – ion channel structure test cases in which peptide toxins at least partially embedded in the membrane environment. Franklin2019 score function is based on REF2015 score terms plus a membrane specific term for computing the implicit membrane environment (Alford et al., 2020).

The available structures of peptide toxin – ion channel complexes for the test cases were optimized using the simple fast relax protocol (Conway et al., 2014) for nuclear magnetic resonance (NMR) and X-Ray structures, and fast relax guided by the experimental electron microscopy (EM) map (Wang et al., 2016) for cryo-electron microscopy (cryoEM) structures (see Methods). We define “near-native” models (NNM) as those which have less than 1 Å root mean square deviation (RMSD) of the C**α**atoms of peptide toxin in each toxin – channel model to its coordinates in the corresponding optimized experimental (native) toxin – channel structure. To explore an unbiased sampling, the peptide toxins were positioned manually in three different initial locations between 3 and 5 Å RMSD from the native binding site. We generated 20,000 models per initial location input, resulting in 60,000 docked models for each test case.

### Membrane energy function improves sampling and scoring for GMTs interacting only with the VSDs

Huwentoxin-IV (HWTX-IV) and Protoxin-II (ProTx-II) from tarantula venom are peptide toxins that have subtype selectivity for and state-dependent antagonist activity against human Na_V_1.7 channel (Schmalhofer et al., 2008; Xiao et al., 2010) that play a key role in the transmission of pain signals (Bennett et al., 2019). HWTX-IV and ProTx-II peptides have been used as scaffolds for optimization of peptides targeting NaV1.7 to develop novel therapeutics to treat pain (Adams et al., 2022; Flinspach et al., 2017; Jiang et al., 2021c; Lawrence et al., 2019; Neff and Wickenden, 2021; Neff et al., 2020). ProTx-II and HWTX-IV inhibit activation of Na_V_1.7 by trapping the VSDII in a deactivated state (IC_50_ ∼ 1 nM for ProTx-II and ∼20 nM for HWTX-IV) (Park et al., 2014; Peng et al., 2002; Schmalhofer et al., 2008; Sokolov et al., 2008; Xiao et al., 2010, 2011; Xu et al., 2019). Although the antagonist effect of these two peptide toxins is achieved by binding to the deactivated states of VSDs, they can also bind to activated states with a lower affinity (Huang et al., 2022; Shen et al., 2019; Wisedchaisri et al., 2020; Xu et al., 2019).

We tested our protein-protein docking protocol on the native structures of ProTx-II bound to a deactivated and activated chimeric hNa_V_1.7-VSDII-Na_V_Ab channel (PDBs: 6N4R and 6N4I, respectively) (Xu et al., 2019) and HWTX-IV bound to a deactivated mutant chimeric hNa_V_1.7-VSDII-Na_V_Ab channel (PDB: 7K48)(Wisedchaisri et al., 2020). We selected 6N4I over 6N4Q structure of activated chimeric hNa_V_1.7-VSDII-Na_V_Ab channel due to the higher resolution of the former. Analysis of all generated ProTx-II -hNa_V_1.7-VSDII-Na_V_Ab channel complex models showed an excellent correlation between the Rosetta interface score and the accuracy of the models (Figure 3), with the top (lowest score values) interface score models having low RMSD values when compared to the native structures (Table 1). Top interface score models for the 6N4I test case were within 0.9 and 0.3 Å RMSD from the native structure using REF2015 or franklin2019, respectively, and for the 6N4R test case were within 0.4 Å RMSD from the native structure using franklin2019 only (Figure 3AB, Table 2). For the 6N4R test case (using REF2015) (Figure 3B i, green cartoon) and for the 7K48 test case (3C i green and blue cartoons), the top interface scoring models were not near-native (> 1.0 Å RMSDs) due to the appearance of a second interface score minimum away from the native binding site. However, when we first extracted the top 10% of models (∼6,000) based on the total Rosetta score (sum of all score terms of the corresponding energy function) and then sorted these models based on the interface score for the 7K48 test case, the top interface score models were within 0.7 and 0.5 Å RMSDs from the native structure using REF2015 and franklin2019, respectively (Figure 3C iv and v, Table 2). Overall, franklin2019 generated a similar or a greater number of NNMs compared to REF2015 and performed better at the scoring of 6N4R, 6N4I, and 7K48 test cases (Tables 1 & 2).

**Table 1.** Results of the benchmarking of our modeling protocol (Part 1)

| Test case | REF2015 |  |  |  |  | franklin2019 |  |  |  |  |
| --- | --- | --- | --- | --- | --- | --- | --- | --- | --- | --- |
|  | Total NNM | top100 I_sc models |  | top100 I_sc (ASBS) models |  | Total NNM | top100 I_sc models |  | top100 I_sc (ASBS) models |  |
|  |  | RMSD (Å) | No. of NNM | RMSD (Å) | No. of NNM |  | RMSD (Å) | No. of NNM | RMSD (Å) | No. of NNM |
| 6N4I | 492 | 1.3 ± 2.0 | 80 | 0.6 ± 0.7 | 85 | 1577 | 0.5 ± 0.3 | 92 | 0.3 ± 0.1 | 100 |
| 6N4R | 157 | 6.1 ± 3.8 | 30 | 5.0 ± 3.7 | 38 | 72 | 2.8 ± 3.2 | 47 | 4.5 ± 1.8 | 7 |
| 7K48 | 32 | 7.2 ± 0.04 | 5 | 6.6 ± 2.2 | 9 | 47 | 5.4 ± 3.0 | 15 | 5.9 ± 2.2 | 12 |
| 6NT4 | 16 | 3.5 ± 0.6 | 0 | 3.7 ± 1.1 | 2 | 84 | 3.3 ± 0.8 | 2 | 3.00 ± 1.7 | 22 |
| 7K18 | 4 | 5.3 ± 0.5 | 0 | 5.1 ± 1.2 | 2 | 40 | 4.9 ± 0.2 | 0 | 4.7 ± 0.5 | 0 |
| 6A95 | 654 | 0.1 ± 0.04 | 100 | 0.1 ± 0.03 | 100 | 430 | 0.5 ± 0.6 | 91 | 1.2 ± 1.2 | 62 |
| 2A9H | 36 | 2.5 ± 1.3 | 21 | 3.6 ± 1.0 | 5 |  |  |  |  |  |
| 4JTA | 1196 | 0.5 ± 0.8 | 80 | 1.6 ± 1.2 | 33 |  |  |  |  |  |
| 7MIX | 0 | 2.4 ± 0.4 | 0 | 3.0 ± 0.4 | 0 |  |  |  |  |  |
| 7MIX* | 2651 | 0.05 ± 0.05 | 100 | 0.04 ± 0.02 | 100 |  |  |  |  |  |
| 6J8E | 0 | 3.0 ± 1.1 | 0 | 2.8 ± 0.9 | 0 |  |  |  |  |  |
| 6J8E* | 624 | 0.04 ± 0.02 | 100 | 0.06 ± 0.08 | 100 |  |  |  |  |  |

This table shows the results of the benchmarking conducted using REF2015 and franklin2019 energy functions for each test case. Total NNM: total number of near native models generated. <RMSD>: average RMSD against native structure of top 100 interface scoring (I\_sc) models; +/- indicated standard deviation. #NNM: number of near native models within the top 100 I\_sc models. ASBS: after score-based selection of top 10% models.
\* For 7MIX and 6J8E, starred rows indicate that the native structure location of the toxin was used as initial location in those benchmarks.

**Table 2.**
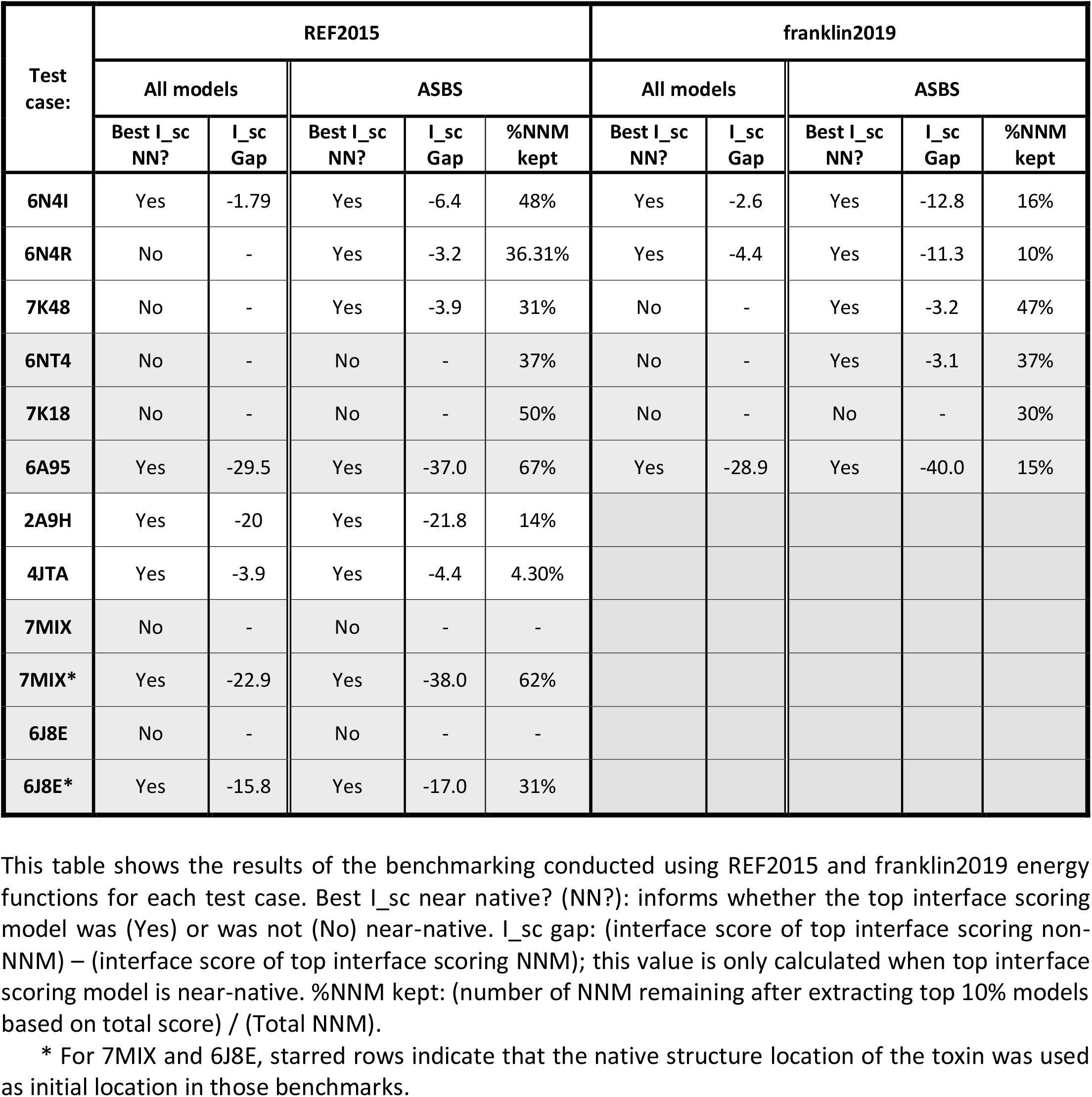
Results of the benchmarking of our modeling protocol (Part 2)

| Test case: | REF2015 |  |  |  |  | franklin2019 |  |  |  |  |
| --- | --- | --- | --- | --- | --- | --- | --- | --- | --- | --- |
|  | All models |  | ASBS |  |  | All models |  | ASBS |  |  |
|  | Best I_sc NN? | I_sc Gap | Best I_sc NN? | I_sc Gap | %NNM kept | Best I_sc NN? | I_sc Gap | Best I_sc NN? | I_sc Gap | %NNM kept |
| 6N4I | Yes | -1.79 | Yes | -6.4 | 48% | Yes | -2.6 | Yes | -12.8 | 16% |
| 6N4R | No | - | Yes | -3.2 | 36.31% | Yes | -4.4 | Yes | -11.3 | 10% |
| 7K48 | No | - | Yes | -3.9 | 31% | No | - | Yes | -3.2 | 47% |
| 6NT4 | No | - | No | - | 37% | No | - | Yes | -3.1 | 37% |
| 7K18 | No | - | No | - | 50% | No | - | No | - | 30% |
| 6A95 | Yes | -29.5 | Yes | -37.0 | 67% | Yes | -28.9 | Yes | -40.0 | 15% |
| 2A9H | Yes | -20 | Yes | -21.8 | 14% |  |  |  |  |  |
| 4JTA | Yes | -3.9 | Yes | -4.4 | 4.30% |  |  |  |  |  |
| 7MIX | No | - | No | - | - |  |  |  |  |  |
| 7MIX* | Yes | -22.9 | Yes | -38.0 | 62% |  |  |  |  |  |
| 6J8E | No | - | No | - | - |  |  |  |  |  |
| 6J8E* | Yes | -15.8 | Yes | -17.0 | 31% |  |  |  |  |  |

This table shows the results of the benchmarking conducted using REF2015 and franklin2019 energy functions for each test case. Best I\_sc near native? (NN?): informs whether the top interface scoring model was (Yes) or was not (No) near-native. I\_sc gap: (interface score of top interface scoring non-NNM) – (interface score of top interface scoring NNM); this value is only calculated when top interface scoring model is near-native. %NNM kept: (number of NNM remaining after extracting top 10% models based on total score) / (Total NNM).
\* For 7MIX and 6J8E, starred rows indicate that the native structure location of the toxin was used as initial location in those benchmarks.

**Figure 3.**
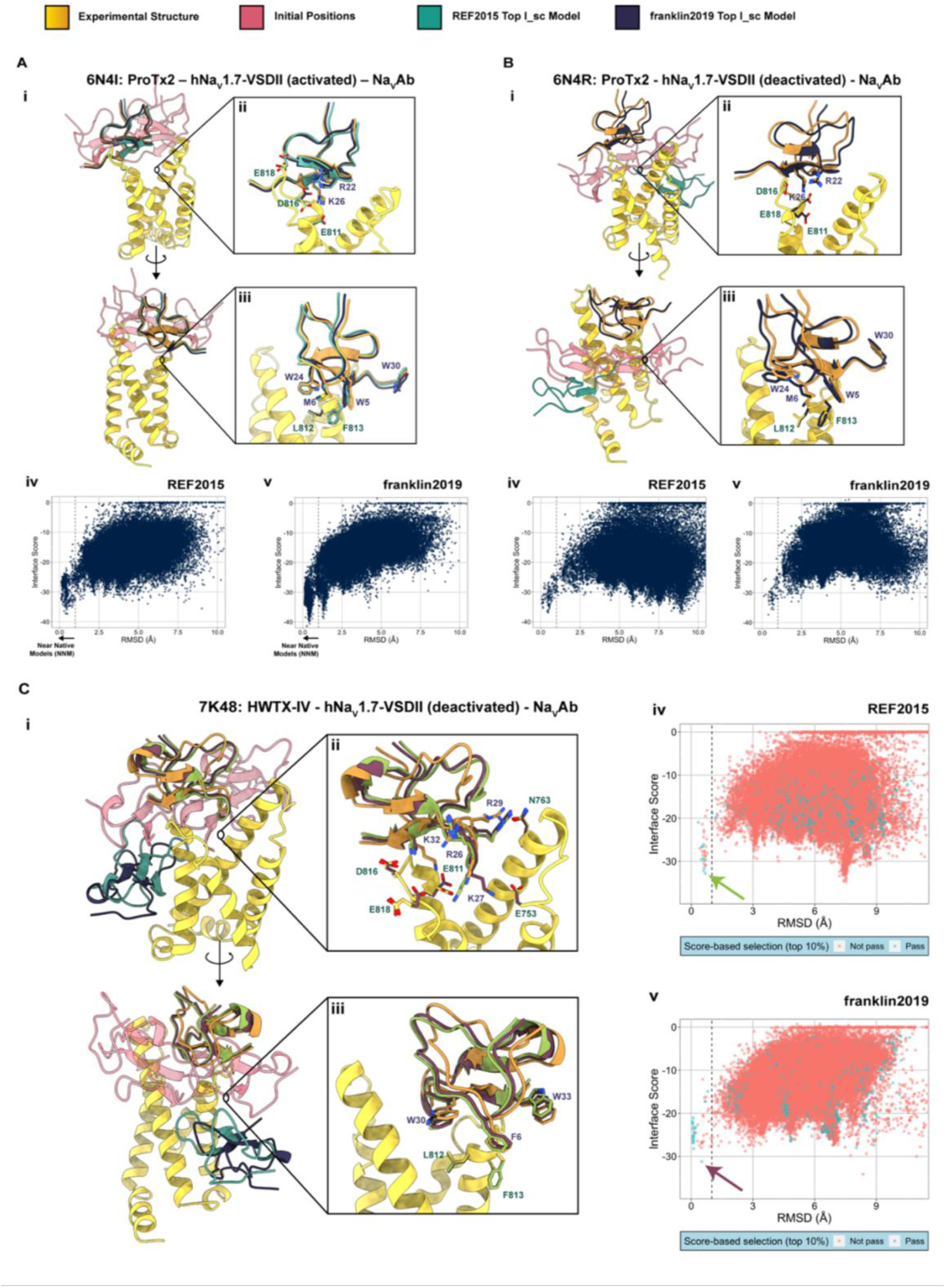
Protein-protein docking of test cases involving gating modifier toxins that only interact with the voltage-sensing domain. (A) Structural modeling of ProTx-II - activated chimeric hNa_V_1.7-VSDII-Na_V_Ab channel complex (native structure PDB: 6N4I) (Xu et al., 2019). i) Front and back view of the channel VSDs with the native structure position of the toxin (yellow), initial locations for RosettaDock (red), and top models from protein – protein docking using REF2015 (green) and franklin2019 (blue). ii) & iii) Front and back detailed views of toxin – channel interfaces. Side-chains of key residues involved in the binding are shown in stick representation and labeled. iv) & v) Plots show RMSD versus interface scores plots for protein – protein docking using REF2015 and franklin2019 for ∼60,000 generated models. (B)Structural modeling of ProTx-II - deactivated chimeric hNa_V_1.7-VSDII-Na_V_Ab channel complex (native structure PDB: 6N4R) (Xu et al., 2019). Subpanels i-v) show data described as in panel A. (C)Structural modeling of HwTx-IV - deactivated mutant chimeric hNa_V_1.7-VSDII-Na_V_Ab channel complex (native structure PDB: 7K48) (Wisedchaisri et al., 2020). Subpanels i-v) show data described as in panel A. The plots in subpanels iv) and v) show data points for the top 10% total score models colored in blue and data points for the rest of the models colored in red.

Subsequently, we visually inspected peptide toxin - channel interfaces in our top interface score models. For the 6N4I test case, the contacts between ProTx-II and the channel at the polar (Figure 3A ii) and membrane (Figure 3A iii) interfaces in our top models are in agreement with the native structure. For the 6N4R test case, ProTx-II presented ∼0.4 Å RMSD backbone shift in the top model compared to the native structure (Figure 3B ii & iii). The top interface score models selected after total Rosetta score-based filtering for the 7K48 test case revealed 0.5-0.7 Å RMSDs backbone shifts of the toxin and other differences in side-chain conformations, particularly at the toxin – channel polar interface (Figure 3C ii). For example, HWTX-IV toxin residue K32 in our top models presents a different rotamer that interacts with channel D816 instead of E811, as is the case in the native 7K48 structure. Another example is HWTX-IV toxin residue R29 which has different side-chain conformations in our top models but maintains the interaction with N763 from the S2 segment of the channel.

### GMTs that interact both with the pore and VSDs might require explicit glycan representation on the pore for accurate modeling

α-Scorpion toxins bind to Na_V_ channels inhibiting fast inactivation and, consequently, alter neuron electrical properties and excitability (Ahern et al., 2016; Bosmans and Tytgat, 2007; Catterall et al., 2007). The structural mechanisms of α-scorpion modulation of Na_V_ channels were recently described (Clairfeuille et al., 2019). α-Scorpion toxins bind to the VSDIV of Na_V_ channels and trap the voltage sensor in the “down” (or deactivated) state. Since activation of Na_V_-VSDIV is required to trigger a fast inactivation (Capes et al., 2013; Jiang et al., 2021b), the α-scorpion toxins bound Na_V_ channels are not able to inactivate, resulting in sustained sodium currents. The receptor site for these toxins, located at the extracellular side of the VSDIV, also comprises interactions with the pore domain I (PDI) residues and glycans (Clairfeuille et al., 2019).

AaH2 is an α-scorpion toxin that inhibits Na_V_1.7-mediated sodium currents (IC_50_ ∼ 50 nM) and its native structure in a complex with a chimeric Na_V_1.7 - Na_V_PaS channel was recently determined (PDB: 6NT4) (Clairfeuille et al., 2019). Our protein-protein modeling protocol sampled NNM for the 6NT4 structure test case (Figure 4A, Table 1). However, top interface scoring models were not near-native and were located in a different binding site, involving only interactions with the VSD (Figure 4A i, model colored in green). For franklin2019, the initial score-based selection of the top 10% models removed the non-native interface score minimum (Figure 4A v), as was observed for the 7K48 structure test case above (Figure 3C iv and v). Despite the absence of explicit glycans during the modeling protocol, the franklin2019 top interface score model revealed an excellent agreement of interactions at the toxin-channel interface (Figure 4A ii & iii) and presented an RMSD of 0.1 Å against the native structure.

**Figure 4.**
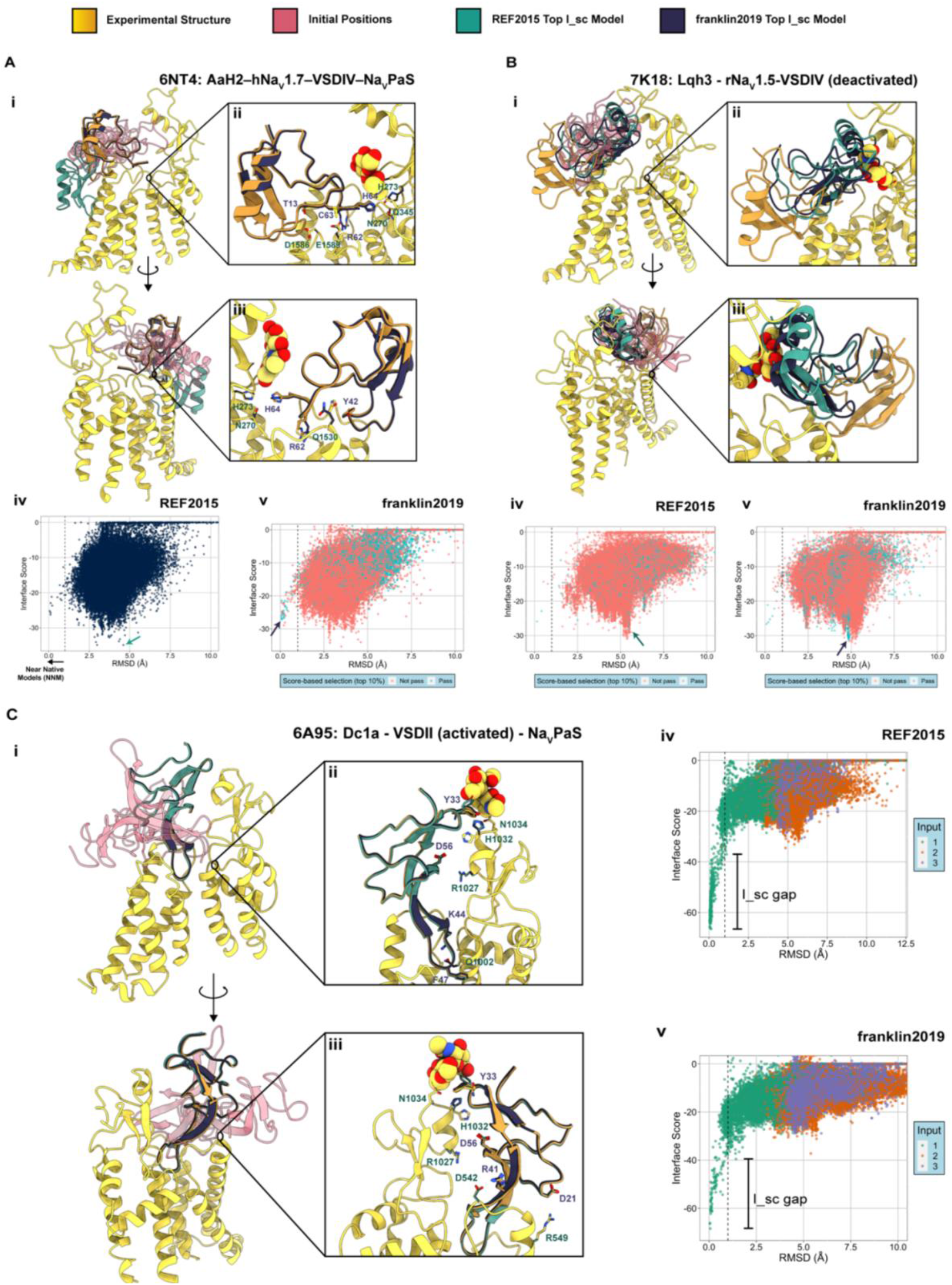
Protein-protein docking of test cases involving gating modifier toxins that interact both with the voltage-sensing and pore domains. (A) Structural modeling of AaH2 - chimeric Na_V_1.7 - Na_V_PaS channel complex (native structure PDB: 6NT4) (Clairfeuille et al., 2019). i) Front and back view of the channel VSDs and PDs with the native structure position of the toxin (yellow), initial locations for RosettaDock (red), and top models from protein – protein docking using REF2015 (green) and franklin2019 (blue). ii) & iii) Front and back detailed views of toxin – channel interfaces. Side-chains of key residues involved in the binding are shown in stick representation and labeled. iv) & v) Plots show RMSD versus interface scores plots for protein – protein docking using REF2015 and franklin2019 for ∼60,000 generated models. The plots in subpanels iv) and v) show data points for the top 10% total score models colored in blue and data points for the rest of the models colored in red. (B) Structural modeling of LqhIII - Na_V_1.5 channel complex (native structure PDB: 7K18) (Jiang et al., 2021a). Subpanels i-v) show data described as in panel A. (C)Structural modeling of Dc1a - Na_V_PaS channel complex (native structure PDB: 6A95) (Shen et al., 2018). Subpanels i-v) show data described as in panel A. The plots in subpanels iv) and v) show data points colored based on the original 3 input positions of Dc1a peptide toxin (green, purple, and orange) that generated each model.

The α-toxin LqhIII from the deathstalker scorpion inhibits fast inactivation of Na_V_1.5 with an IC_50_ ∼ 10 nM and the cryoEM structure of this toxin bound to the cardiac sodium channel was recently resolved (Jiang et al., 2021a) (PDB: 7K18). RosettaDock showed similar results for 7K18 as for the 6NT4 structure test case above, with a sampling of NNMs but and the appearance of a non-native interface score minimum (Figure 4B iv & v). Moreover, after score-based selection, top interface scoring models were still in the non-native minimum for both REF2015 and franklin2019. In these models, LqhIII is located in a binding site occupied by glycans in the native structure (Figure 4B ii & iii), suggesting that explicit glycan representation might be necessary to potentially generate top interface scoring models near the native structure.

Dc1a is a spider toxin that binds to activated VSDII from insect Na_V_s promoting channel opening (Bende et al., 2014). We used the cryoEM structure of Dc1a bound to Na_V_PaS (PDB: 6A95) (Shen et al., 2018) as another test case for GMTs with interactions at the voltage-sensor and pore domains. In contrast to the two previous cases, our modeling protocol was able to sample NNMs and converge on interface score minimum with a large interface score gap compared to non-native models (Figure 4C iv & v). Top interface scoring models from REF2015 and franklin2019 benchmarks both where within 0.1 Å RMSDs from the native structure. Interestingly, most NNM were generated from the same input initial location (Figure 4C iv & v, green data points; Figure S1AB), while one of the other inputs did not generate any NNM (Figure 4C iv & v, purple data points; Figure S1AB). The binding interfaces of top models were essentially identical to the native structure despite comprising interactions with glycans, which were not present during the modeling (Figure 4C i, ii & iii).

### Spatial constraints may hamper sampling of peptide toxin – ion channel interactions at the pore domain

Charybdotoxin (CTX) is a pore-blocking toxin of K^+^ channels from scorpion venom (Garcia et al., 1995; Miller, 1995; Miller et al., 1985). We tested our protein-protein docking protocol on the X-ray structure of CTX - rK_V_1.2-2.1 chimera (PDB: 4JTA) (Banerjee et al., 2013). Interestingly, top interface scoring models had RMSDs values around 2.3 Å from the native structure. When we visually analyzed these top models, we found that the toxin was rotated 180° with respect to the native structure (Figure 5A ii), due to the K_V_ channel 4-fold symmetry in the rK_V_1.2-2.1 chimera structure (Banerjee et al., 2013). We rotated the native structure of the channel 180° around the Z-axis and then recalculated the RMSDs, obtaining RMSD values ∼0.5 Å for top interface scoring models (Figure S2), showing that these models are near the native 4JTA structure. Despite the rotated pose of CTX toxin relative to the channel pore, critical CTX residue for the pore-blocking effect (K27) positioned similarly in our top interface score model and the native structure (Figure 5A ii).

**Figure 5.**
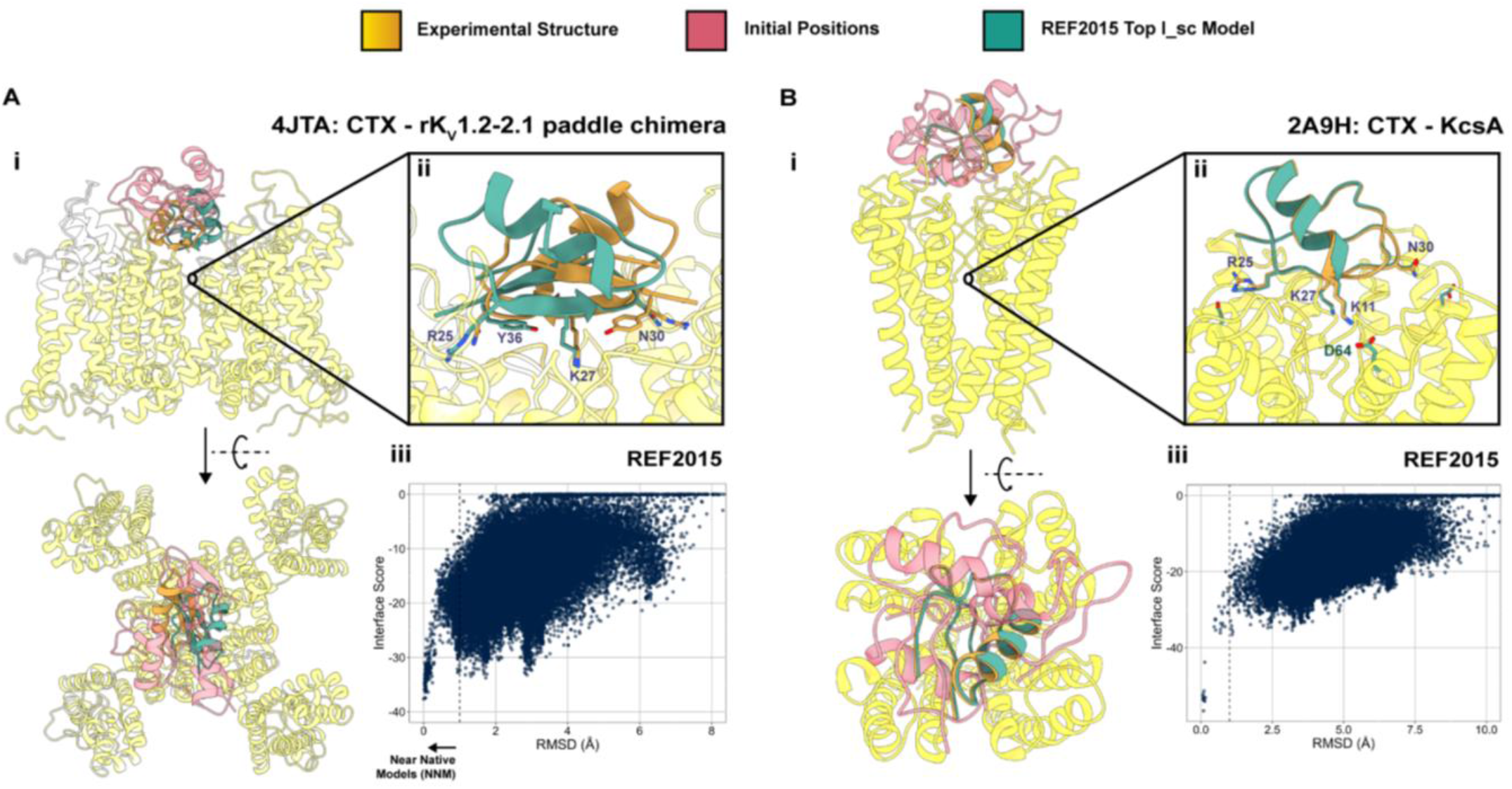
Protein-protein docking of test cases involving pore-blocking charybdotoxin (CTX) and potassium channels. (A) Structural modeling of Charybdotoxin (CTX) – chimeric rK_V_1.2-2.1 channel complex (native structure PDB: 4JTA) (Banerjee et al., 2013). i) Front and outside view of the channel with the native structure position of the toxin (yellow), initial locations for RosettaDock (red), and top model from REF2015 (green). ii) Detailed view of toxin – channel interfaces. Side-chains of key residues involved in the binding are shown in stick representation and labeled. iii) Plots show RMSD versus interface scores plots for protein – protein docking using REF2015 for ∼60,000 generated models. (B) Structural modeling of CTX - KcsA channel complex (native structure PDB: 2A9H) (Yu et al., 2005). Subpanels i-iii) show data described as in panel A.

The structure of CTX in complex with the K^+^ channel KcsA is the only NMR structure in our peptide toxin – ion channel benchmark (PDB: 2A9H) (Yu et al., 2005). Notably, the outer vestibule of KcsA was mutated to bind CTX in this structure. Our protein-protein docking protocol generated relatively small number of NNMs (Figure 5B iii, Table 1), with all of them originating from the same input initial location, highlighting again how the initial location of the peptide biased the outcome of the protein-protein docking (Figure S1C). Our top interface score models were within 0.1 Å RMSD from the native structure and adequately captured interactions at the protein-protein interface (Figure 5B ii).

Conotoxins are a group of neurotoxic peptides isolated from the venom of cone snails that target and block ion channels with high selectivity and affinity (Green and Olivera, 2016; Terlau and Olivera, 2004). Our original protein-protein docking protocol was unable to generate NNMs for the pore blocking conotoxin ziconotide bound to hCa_V_2.2 (PDB: 7MIX) (Gao et al., 2021) and for the pore blocking conotoxin KIIIA bound to hNa_V_1.2 (PDB: 6J8E) (Pan et al., 2019) (Figure 6AB iii, Table 1). For the 7MIX structure test case, the top interface scoring model was located in the same binding site as observed in the native structure but turned upside down (Figure 6A, model colored in green). In the 6J8E structure test case, the top interface scoring model was located outside of the channel pore, despite using initial positions within the pore (Figure 6B, model colored in green). To evaluate if this was a problem of sampling or scoring, we re-ran our protocol for these two test cases using the native pose as the initial input location to bias the sampling within the native binding site. For both the 7MIX and 6J8E structure test cases, our protein-protein docking protocol generated convergent interface score funnels when using the native pose as an input location (Figure 6AB iii, pink dots), which highlights the ability of the Rosetta scoring function to distinguish between near native and non-native models. Notably, even when using the native pose as the initial location in the 7MIX and 6J8E test cases, RMSD – interface score plots reveal a lack of sampling ∼1 Å RMSD from the native structure position of peptide toxin (Figure 6AB iii, dashed line) and suggesting a sampling challenge in these test cases. This result reveals examples of large energetic barriers to reaching the binding site for these peptide toxins that is caused by the spatial structural features within the extracellular region of the pore. Both conotoxins bind to small and constrained binding sites surrounded by the long extracellular loop regions. Therefore, extensive sampling of peptide toxin poses within the constrained binding sites might be necessary to potentially generate top interface scoring models near the native structure for peptide toxins targeting binding sites surrounded by the long extracellular loop regions of ion channels.

**Figure 6.**
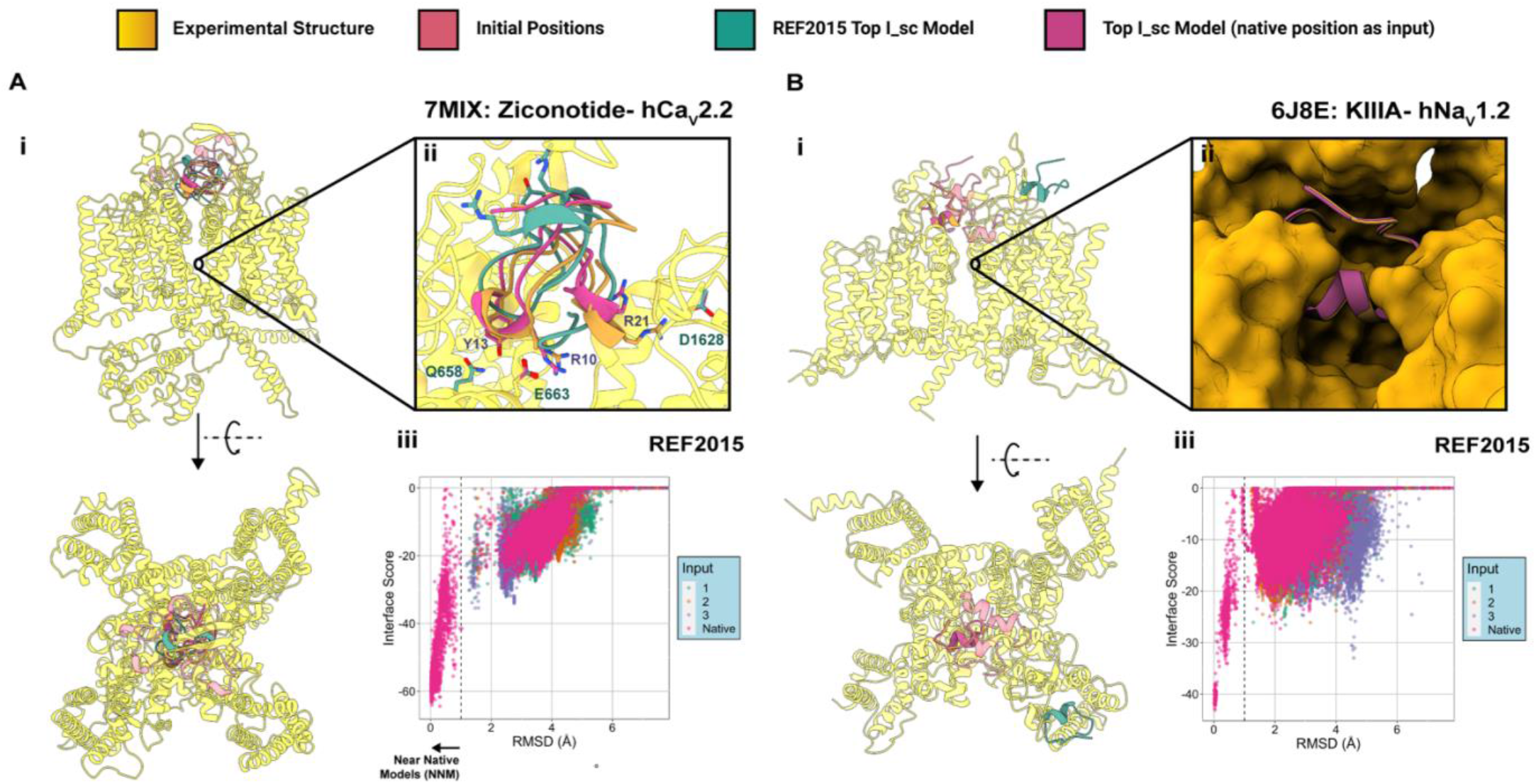
Protein-protein docking of test cases involving pore-blocking conotoxins and Nav and Cav channels. (A) Structural modeling of conotoxin ziconotide - hCa_V_2.2 channel complex (native structure PDB: 7MIX) (Gao et al., 2021). i) Front and outside view of the channel with the native structure position of the toxin (yellow), initial locations for RosettaDock (red), and top model from REF2015 (green). ii) Detailed view of toxin binding site. For 6J8E the molecular surfaces are shown to highlight the spatial constraints of the toxin binding site. iii) Plots show RMSD versus interface scores plots for protein – protein docking using REF2015 for ∼80,000 generated models (original 60,000 plus 20,000 using the native position as input). (B) Structural modeling of conotoxin KIIIA – hNav1.7 channel complex (native structure PDB: 6J8E) (Pan et al., 2019). Subpanels i-iii) show data described as in panel A.

## DISCUSSION

Peptide toxins have been essential molecular tools for studying the function, structure, and dynamics of ion channels for decades (Dutertre and Lewis, 2010; Herzig et al., 2020; Norton and Chandy, 2017). Moreover, they are promising biomolecules to develop biologics as therapeutics to target and modulate specific ion channels with superior subtype and state selectivity (Bordon et al., 2020; Neff and Wickenden, 2021; Nguyen and Yarov-Yarovoy, 2022; Norton, 2017; Norton and Chandy, 2017; Wulff et al., 2019). Limitations in obtaining high-resolution structural data to understand and harness the diverse molecular mechanisms of peptide toxins action on ion channels hamper the drug development process to optimize and translate these peptides to the clinic.

Computational structural modeling has emerged as a powerful tool to advance research in the ion channel field (Craig et al., 2020; Gupta et al., 2015; Pumroy et al., 2022; Tilley et al., 2014; Tuluc et al., 2016; Vargas et al., 2012; Yang et al., 2015, 2018; Yarov-Yarovoy et al., 2012; Zhang et al., 2012). For example, homology modeling is a useful approach for modeling ion channels lacking high-resolution structures (Doupnik et al., 2015; Tang et al., 2017; Tikhonov and Zhorov, 2014). Additionally, novel protein structure prediction algorithms based on deep learning, AlphaFold and RoseTTAFold (Baek et al., 2021; Jumper et al., 2021), enable ion channel structure prediction in one particular state with relatively high accuracy, although *de novo* modeling of specific ion channel states remains challenging. Ligand docking has been used to map the binding sites of small molecules in ion channels and get insights into molecular mechanisms of drug action (Du et al., 2014; Madeja et al., 2010; Melgari et al., 2015; Nguyen et al., 2017; Tikhonov and Zhorov, 2014). In addition, both structural models and native structures represent only snapshots of the full spectrum of protein dynamics. Using static ion channel structures as inputs for molecular dynamics simulations can provide a broader perspective of the dynamics underlying the ion channel gating and modulation (Flood et al., 2019; Galleano et al., 2021; Nguyen et al., 2019; Rovšnik et al., 2021; Suma et al., 2020). Finally, structure-guided computational protein design has achieved significant milestones in recent years (Anishchenko et al., 2021; Cao et al., 2020, 2022; Sahtoe et al., 2022; Silva et al., 2019), and these approaches can be applied to engineer ion channels (Xu et al., 2020; Yang et al., 2016) or design novel ion channel modulators (Xu et al., 2021). In this study, we developed a computational protein-protein docking protocol using Rosetta to model the interactions between venom peptide toxins and ion channels. We tested our protocol on available native structures of peptide toxin – ion channel complexes representing a diversity of venom peptides and ion channel subfamilies. Detailed analysis of each of the test cases revealed the strengths and limitations of our protein-protein docking approach.

We confirmed that the Rosetta score term that best correlates with model accuracy is the interface score (Chaudhury et al., 2011). For the standard Rosetta energy function (REF2015), in 4 out of 10 test cases, the model with the top interface score was near-native (Table 2). For the Rosetta membrane energy function (franklin2019) that was tested only on peptide toxins at least partially embedded in the membrane environment, in 3 out of 6 test cases, the model with the top interface score was near-native (Table 2). However, when we first extracted the top 10% models based on total Rosetta score and then focused on the models with the top interface score, we identified NNMs in 6 out of 10 test cases for REF2015, and identified NNMs in 5 out of 6 test cases for franklin2019 (Table 2). Therefore, using the total Rosetta score as an initial filter improved the identification of an NNM as the top interface scoring model. This improvement can be explained by looking at the RMSD – interface score plots of the 7K48 & 6NT4 test cases (Figure 3C; Figure 4A), where we see the appearance of non-native interface score minima populated with models that are excluded after extracting the top 10% scoring models. This result highlights the importance of identifying models with predicted strong binding (interface score) and realistic energy values (total Rosetta score). Notably, score-based selection did not allow us to remove the non-native interface score minimum for 7K18 (Figure 4B), suggesting that explicit glycan representation might be necessary to potentially generate top interface scoring models near the native structure.

Experimental data highlights the importance of membrane embedding in the function of some of the peptide toxins tested in our study, particularly for GMTs (Agwa et al., 2017; Henriques et al., 2016; Lee and MacKinnon, 2004; Mihailescu et al., 2014; Mueller et al., 2020; Zhang et al., 2019). We took advantage of Rosetta’s versatility and implemented the latest membrane energy function (franklin2019) in our modeling protocol to evaluate its performance compared to the standard energy function (REF2015). In the 6 test cases tested with both energy functions, we observed that franklin2019 performs similarly to or better than REF2015 in generating NNMs and more accurate models based on average RMSD of top scoring models (Table 1). The effect of implicit accounting of membrane environment in scoring can be visually identified if we compare the RMSD – interface score plots of franklin2019 and REF2015 for the 6N4I and 6N4R test cases (Figure 3AB). The RMSD – interface score plots for franklin2019 are able to maintain a funnel-like shape, while RMSD – interface score plots for REF2015 have a shallow slope shape due to the appearance of models with low interface score values with relatively large RMSDs values. These non-native REF2015 energy function based models are actually placed within the membrane environment (Figure 3B, green cartoon); confirming that scoring peptide toxins that are at least partially embedded in the membrane environment with franklin2019 energy function is necessary to generate more NNMs.

Our benchmarking study also highlighted the limitations of our current approach. Since we are using a Monte Carlo – based algorithm, the protocol may not be able to thoroughly sample near-native models and the scoring may not be able to identify near-native models from non-native models. For the α - scorpion toxins AaH2 and Lqh3, we observed that while our protein-protein docking approach can generate NNMs, these models are not representing in the lowest interface score minimum (Figure 4AB). Notably, the α -scorpion toxins are known to interact with glycans, the result of post-translational modifications of the ion channel (Clairfeuille et al., 2019; Jiang et al., 2021a) and glycans were not accounted for in the current version of our protocol. If we look at the position of Lqh3 with respect to the channel in the non-native lowest interface score minimum, we see that the actual placement of the α -scorpion toxins would clash with glycan moieties if they were present (Figure 4B i & iii, green and blue cartoons). This suggests that explicit placement of glycans would remove this non-native interface score minimum. Glycans will be implemented and tested in future updates of our protocol.

Finally, we were not able to generate NNMs for 7MIX and 6J8E test cases, which correspond to the conotoxins ziconotide and KIIIA respectively (Table 1). To understand whether near-native conformations were being sampled and rejected due to scoring problems or not being sampled, we reran our modeling protocol using the native structure position as the initial location of the toxin to bias the Monte Carlo search around the native binding site. Indeed, this bias allowed us to generate NNMs that sampled deep interface score minima in both cases (Figure 6A iii; 6B iii). This result indicates that the original limitation of our protein-protein docking approach was due to a sampling problem and not a scoring problem, as was the case for α-scorpion toxins. To understand why Rosetta struggled with sampling near-native conformations for these two specific test cases, we looked at the toxin binding sites. Both toxins are pore blockers and bind within relatively small and constrained binding sites where the channel’s extracellular long loop regions surround the toxin (Figure 6A i; Figure 6B i & ii), in contrast to GMTs or CTX which can relatively freely move and rotate near their binding sites without clashing with the channel. This means that most of the Monte Carlo perturbations within relatively small and constrained binding sites will be rejected due to steric clashes, which limits progression through the sampling. Using more than 3 initial locations of a toxin or using smaller perturbations during the Monte Carlo search are possible ways to address the sampling limitations. Notably, there were multiple test cases in which some initial locations generated 0 NNMs, or where all NNMs came from the same original input (Figure S1). This highlights the importance of using multiple initial locations of a toxin to increase the chances of sampling near native conformations.

Visual inspection of peptide toxin – ion channel interfaces of NNMs, showed a recapitulation of specific protein – protein interactions in all test cases. Our models revealed that in some test cases, side-chain rotamers at the protein – protein interface were identical between our models and native structures (Figure 3A ii & iii; Figure 4C ii & iii), and in other test cases side-chain rotamers at the protein – protein interface were different but maintained key interactions (Figure 3C ii & iii; Figure 5A ii & iii). Notably, in some test cases, NNMs presented small shifts in toxin backbone coordinates compared to the native structure, as is the HWTX-IV case (Figure 3C). Despite these backbone shifts, interactions at toxin – ion channel interfaces are, in most cases, captured. These differences among our models and native structures might represent alternative binding poses within the same energy minimum.

Overall, our modeling protocol and results of test cases present a proof of concept and point of reference for future projects that require understanding the molecular determinants and mechanisms of peptide toxin interactions with ion channels. We anticipate that as the accuracy of computational structural modeling approaches increases, protein – protein docking of peptide – ion channel complexes will be useful for design and optimization of peptide-based therapeutics targeting ion channels.

## METHODS

### 1. RESOURCE AVAILABILITY

#### Lead contact

For more information and requests for specific codes or computational models, please contact Vladimir Yarov-Yarovoy.

#### Materials availability

NA

#### Data and code availability

To use this protocol, the Rosetta software package is required. This package is free for academic users, and information on licensing can be found at www.rosettacommons.org. The software package should be used that is dated after March, 2020. Specific code and execution options have been provided in Methods.

Any additional information required to reanalyze the data reported in this paper is available from the lead contact upon request.

### 2. EXPERIMENTAL MODEL AND SUBJECT DETAILS

NA

### 3. METHOD DETAILS

#### Molecular graphics visualization

All models and structures were processed and analyzed using UCSF ChimeraX (Goddard et al., 2018).

##### Structure optimization

Native structures were downloaded from the RCSB PDB (https://www.rcsb.org/). Firstly, the structures were cleaned:

~~~
*$ROSETTAPATH/main/tools/protein_tools/scripts/clean_pdb.py$pdb $chainsID*
~~~

For cryoEM structures, optimization was guided by the EM map (Conway et al., 2014; Wang et al., 2016). This requires the following RosettaScripts (Bender et al., 2016) XML protocol:

~~~
*<ROSETTASCRIPTS>
<SCOREFXNS>
 <ScoreFunction name=“dens” weights=“ref2015”>
  <Reweight scoretype=“elec_dens_fast” weight=“35.0”/>
<Set
scale_sc_dens_byres=“R:0.76,K:0.76,E:0.76,D:0.76,M:0.76,C:0.81,Q:0.81,H:0.81,N:0.81,T:0.81,S:0.81,Y: 0.88,W:0.88,A:0.88,F:0.88,P:0.88,I:0.88,L:0.88,V:0.88”/>
 <Reweight scoretype=“cart_bonded” weight=“0.5”/>
 <Reweight scoretype=“pro_close” weight=“0.0”/>
</ScoreFunction>
<ScoreFunction name=“r15” weights=“ref2015”>
 <Reweight scoretype=“cart_bonded” weight=“0.5”/>
 <Reweight scoretype=“pro_close” weight=“0.0”/>
</ScoreFunction>
</SCOREFXNS>
<MOVERS>
<SetupForDensityScoring name=“setupdens”/>
<LoadDensityMap name=“loaddens” mapfile=“%%map%%”/>
<FastRelax name=“relaxcart” scorefxn=“dens” repeats=“2” cartesian=“1”/>
</MOVERS>
<PROTOCOLS>
<Add mover=“setupdens”/>
<Add mover=“loaddens”/>
<Add mover=“relaxcart”/>
</PROTOCOLS>
<OUTPUT scorefxn=“r15”/>
</ROSETTASCRIPTS>*
~~~

This protocol (named B_relax_density.xml) is run by calling RosettaScripts, together with the native structure, the EM map and map resolution:

~~~
*$ROSETTAPATH/main/source/bin/rosetta_scripts.linuxgccrelease \
-in:path:database $ROSETTAPATH /main/database \
-in:file:s ./$pdb \
-in:file:native ./$pdb \
-parser:protocol ./B_relax_density.xml \
-parser::script_vars map=$map \
-relax:constrain_relax_to_start_coords \
-ignore_unrecognized_res \
-edensity::mapreso $mapres \
-edensity::cryoem_scatterers \
-crystal_refine \
-nstruct $nstruct \
-default_max_cycles 200 \
-out:prefix EM-relax-density- \
-out:file:silent ./outfiles/EM-relax-density-$pdb_${SLURM_ARRAY_TASK_ID}.silent \
-out:file:silent_struct_type binary \
-out:file:scorefile ./outfiles/EM-relax-density-$pdb.sc \*
~~~

For X-ray or NMR structures, optimization was performed similarly, using the FastRelax mover in RosettaScriptts and changing the energy function to REF2015 set for non-ideal minimization (r15), and removing EM-related options.

#### Setting of membrane geometry

To transform the structures into membrane coordinates, they were superimposed into their respective structures downloaded from the PDBTM database (http://pdbtm.enzim.hu/) using the MatchMaker tool of ChimeraX. Note that the reasons for not using the structures from PDBTM directly were i) for consistency due to small disparities observed between these two databases; ii) to be able to use the EM maps to guide initial optimizations.

Cleaned, optimized, and transformed structures were used to generate spanfiles (.span) which define transmembrane regions:

~~~
*$ROSETTAPATH/main/source/bin/mp_span_from_pdb.linuxgccrelease
-in:file:s $pdb \
-ignore_unrecognized_res*
~~~

#### Input preparation

For each test case, the corresponding toxin was placed manually at various positions around the binding site using ChimeraX. Three final inputs with RMSDs between 3 and 5 Å against the optimized native structure were chosen for subsequent steps. Note that these inputs were the same for both REF2015 and franklin2019 tests.

#### Prepacking

Prepacking is required before running docking to optimize side chains outside the interface.

~~~
*$ROSETTAPATH /main/source/bin/docking_prepack_protocol.linuxgccrelease \
-in:file:s ./input.pdb \
-score:weights franklin2019.wts \ # Or ref2015.wts
-nstruct 20 \
-mp:setup:spanfiles input.span \
-mp:scoring:hbond \
-packing:pack_missing_sidechains 0 \
-partners A_B \
-ex1 \
-ex2 \
-docking:sc_min 1 \
-out:suffix _prepack \*
~~~

#### RosettaDock

20,000 docked models per input were generated with RosettaDock using the following options:

~~~
*$ROSETTAPATH/main/source/bin/mp_dock.linuxgccrelease \ # Or docking_protocol.linuxgccrelease
-in:file:s input.pdb \
-in:path:database $ROSETTAPATH/main/database \
-score:weights franklin2019.wts \ # Or ref2015.wts
-mp:setup:spanfiles input.span \
-docking:partners A_B \
-docking:dock_pert 4 10 \
-packing:pack_missing_sidechains 0 \
-nstruct $nstr \
-use_input_sc \
-docking:sc_min 1 \
-ex1 \
-ex2 \
-ex3 \
-ex4 \
-ex1aro \
-ex2aro \
-out:suffix _local_dock \
-out:file:scorefile ./outfiles/LocalDock.fasc \
-out:file:silent ./outfiles/LocalDock- ${SLURM_ARRAY_TASK_ID}.silent \
-out:file:silent_struct_type binary \
-out:file:fullatom \
-score:docking_interface_score 1*
~~~

#### Analysis of docked models

Manipulation of resulting silent files (merging, extracting top scoring subsets, etc.) was conducted with homemade bash scripts and Rosetta scoring application (score_jd2). Analysis of scorefiles (containing total scores, interface scores, and RMSDs against native optimized structures) was performed using homemade RStudio scripts.

### QUANTIFICATION AND STATISTICAL ANALYSIS

NA

### KEY RESOURCES TABLE

5. KEY RESOURCES TABLE
| REAGENT or RESOURCE | SOURCE | IDENTIFIER |
| --- | --- | --- |
| Software and algorithms |  |  |
| Rosetta | Rosetta Commons | https://www.rosettacommons.org/ |
| RosettaDock | NA | https://www.rosettacommons.org/docs/latest/application_documentation/docking/docking-protocol |
| R (Rstudio) | R-project | https://www.r-project.org/<br>https://www.rstudio.com/ |
| ChimeraX | UCSF | https://www.cgl.ucsf.edu/chimerax/ |

## SUPPLEMENTAL INFORMATION

Supplemental Information includes Figures S1 and S2.

## ACKNOWLEDGMENTS

We thank Drs. Heike Wulff and Jon T. Sack and members of the V.Y.-Y. and J.T.S. laboratories for helpful discussions.

## AUTHOR CONTRIBUTIONS

Conceptualization, D.M.L., and V.Y.Y.; Software, D.M.L.; Validation, D.M.L. Resources, D.M.L; Investigation, D.M.L., and V.Y.Y.; Writing – Original Draft, D.M.L.; Writing – Review & Editing, D.M.L., and V.Y.Y.; Supervision, V.Y.Y.

