## Supplemental Information for "Structural Modeling of Peptide Toxin - Ion Channel Interactions using RosettaDock"

**Figure S1**

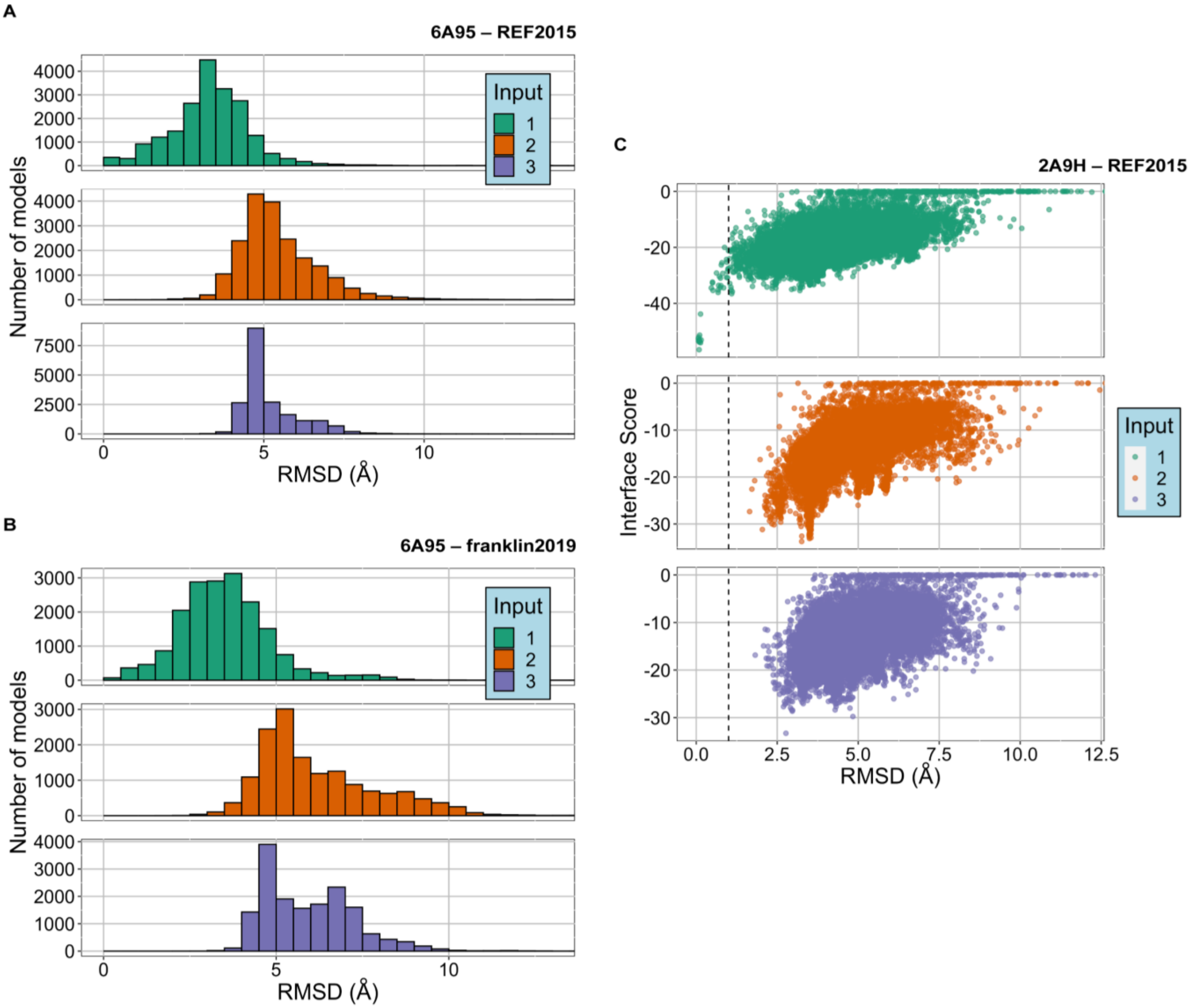


**Figure S1. Initial placement of the toxin biases the outcome of docking simulation**

(A) Distribution of RMSDs of docked models against native structure generated for the 6A95 test case using REF2015. Distributions are divided based on the original input that generated the docked models in order to illustrate that each input resulted in distinct distributions.

(B) Distribution of RMSDs of docked models against native structure generated for the 6A95 test case using franklin 2019.

(C) RMSD – interface score plots of each input for the 2A9H test case.

**Figure S2**

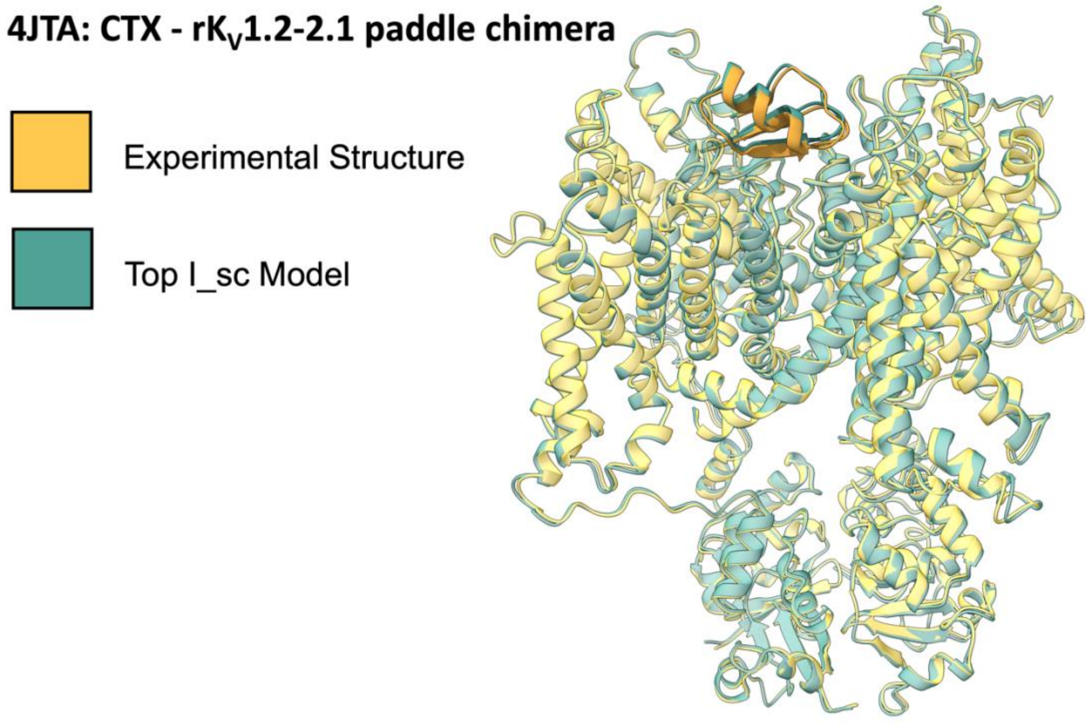


**Figure S2. CTX binding to rK$_V$1.2-2.1 chimera**

The experimental structure of CTX – rK$_V$1.2-2.1 chimera complex (yellow) (PDB ID: 4JTA (Banerjee et al., 2013)) rotated 180˚ around the Z-axis compared to the top interface scoring model from the docking protocol (green) with no rotation.
